# HemaCisDB: An Interactive Database for Analyzing Cis-Regulatory Elements Across Hematopoietic Malignancies

**DOI:** 10.1101/2024.03.12.583848

**Authors:** Xinping Cai, Qianru Zhang, Bolin Liu, Lu Sun, Yuxuan Liu

## Abstract

Noncoding cis-regulatory elements (CREs), such as transcriptional enhancers, are key regulators of gene expression programs. Accessible chromatin and H3K27ac are well-recognized markers for CREs associated with their biological function. Deregulation of CREs is commonly found in hematopoietic malignancies yet the extent to which CRE dysfunction contributes to pathophysiology remains incompletely understood. Here, we developed HemaCisDB, an interactive, comprehensive, and centralized online resource for CRE characterization across hematopoietic malignancies, serving as a useful resource for investigating the pathological roles of CREs in blood disorders. Currently, we collected 922 ATAC-seq, 190 DNase-seq, and 531 H3K27ac ChIP-seq datasets from patient samples and cell lines across different myeloid and lymphoid neoplasms. HemaCisDB provides comprehensive quality control metrics to assess ATAC-seq, DNase-seq, and H3K27ac ChIP-seq data quality. The analytic modules in HemaCisDB include transcription factor (TF) footprinting inference, super-enhancer identification, and core transcriptional regulatory circuitry analysis. Moreover, HemaCisDB also enables the study of TF binding dynamics by comparing TF footprints across different disease types or conditions via web-based interactive analysis. Together, HemaCisDB provides an interactive platform for CRE characterization to facilitate mechanistic studies of transcriptional regulation in hematopoietic malignancies. HemaCisDB is available at https://hemacisdb.chinablood.com.cn/.

## Introduction

Mammalian gene expression program requires coordinated regulation of cis-regulatory elements (CREs), such as promoters, enhancers, silencers, and insulators. A promoter contains DNA sequences located immediately upstream and downstream of a gene to initiate gene transcription [1]. Enhancers are cis-regulatory DNA sequences which can loop over long genomic distances to interact with the promoters of target genes and activate gene expression in a distance-, orientation-, and position-independent manner [2]. Enhancers can be classified into three states, namely, inactive, poised, or active, based on their activity. An inactive enhancer is typically characterized by well-organized unmodified nucleosomes that obstruct the recruitment of transcription factors (TFs) and RNA polymerase. Poised enhancers, which are primed for activation, are often subject to mono-methylation of H3K4 (H3K4me1) modification, which makes chromatin loose and more accessible. In contrast, fully activated enhancers, typically marked by H3K27 acetylation (H3K27ac) modification, exhibit fully open chromatin, facilitating the accessibility of RNA polymerase for bi-directional transcription initiation [3, 4]. Silencers are non-coding genomic sequences which bind repressor proteins and prevent the transcription of target genes [5]. An insulator is a type of CRE that can mediate intra- and inter-chromosomal interactions [6]. CREs contain DNA motifs that are bound by sequence-specific DNA-binding proteins which in turn recruit co-factors to regulate target gene expression [1, 7]. H3K27ac and chromatin accessibility detected by DNase I hypersensitivity or the assay of transposase accessible chromatin with sequencing (ATAC-seq) are commonly used to identify candidate CREs, including promoters and active enhancers, which are the main focus of this study [7, 8].

Emerging evidence shows that deregulation of CREs often contributes to the pathogenesis of hematopoietic malignancies. In T-cell acute lymphoblastic leukemias (T-ALLs), recurrent non-coding mutations that introduce de novo binding sites of MYB, a TF and proto-oncogene, were found to initiate aberrant enhancer complexes and drive the expression of downstream oncogenes including *TAL1* and *LMO1* [9, 10]. Another study with the analysis of 160 T-ALL cases identified recurrent focal duplications on chromosome 8q24 encompassing a NOTCH1-associated enhancer that regulates *MYC* expression [11]. In chronic lymphocytic leukemia (CLL) patients, a densely mutated intergenic region with enhancer characteristics was identified and its disruption causes reduced expression of the B-cell-specific TF *PAX5* [12]. In acute myeloid leukemia (AML) patients with inv(3)/t(3;3), a *GATA2* enhancer rearrangement causes *GATA2* haploinsufficiency and *EVI1* activation simultaneously [13]. It was also found in leukemia cells that noncoding variants at *KRAS* and *PER2* enhancers affect the binding of nuclear receptor family TFs PPARG and RXRA, which disturbs the expression of *KRAS* and *PER2* and promotes the growth of leukemia cells [14]. Moreover, a recent study in diffuse large B cell lymphoma (DLBCL) showed that active super-enhancers (SEs) harboring hypermutation link to the upregulation of oncogenes related to B cell developmental and malignant transformation, including *BCL6*, *BCL2*, and *CXCR4* [15].

CRE landscape also serves as a useful biomarker to identify various subtypes of blood cancers and provides important information about potential cell-of-origin and/or targetable vulnerabilities [16–18]. Enhancer profiling in AML identified an active SE at the gene locus of *RARA* in 25% of samples, and its presence is associated with RARA-directed therapy responses [16]. Thus, the comprehensive and unbiased analysis of CREs is necessary to identify possible genetic and epigenetic causes of diseases, dissect regulatory mechanisms controlling disease initiation and progression, and stratify patients into subgroups for potential personalized medical care. Several databases have been developed to profile enhancer architecture and core transcriptional regulatory circuitry (CRC) in pan-cancer. These databases include Cistrome, ATACdb, dbSUPER, SEA, SEdb, and Cancer CRC [19–24]. Cistrome, a comprehensive database containing DNase-seq, ATAC-seq, and ChIP-seq data for over 1500 TFs and histone marks, lacks disease-specific inquiries, making it challenging for users to locate datasets related to different types of hematopoietic malignancies [19]. Additionally, the Cistrome Cancer module, which is designed for enhancer/SE analyses, is only available for solid tumors. ATACdb, a repository of ATAC-seq data, profiles open chromatin regions across multiple cancer types, but has only a limited number of datasets related to hematopoietic malignancies [20]. The SEA, dbSUPER, and SEdb databases, curating SEs derived from H3K27ac ChIP-seq data across diverse cell types and tissues in various species, also contain a restricted number of datasets relevant to blood cancers [21–23]. Likewise, Cancer CRC, characterizing CRC through the integration of H3K27ac ChIP-seq and ATAC-seq data, solely confines to solid tumors [24]. Therefore, a comprehensive, interactive, and centralized database that characterizes CREs in blood cancers is necessary but currently unavailable.

Here, we profiled CREs across myeloid and lymphoid neoplasms and developed a CRE database HemaCisDB (https://hemacisdb.chinablood.com.cn/) specific to hematopoietic malignancies. We collected 922 ATAC-seq, 190 DNase-seq, and 531 H3K27ac ChIP-seq data from cell lines and patient samples covering the full spectrum of leukemia and lymphoma subtypes, including Hodgkin’s lymphoma (HL), non-Hodgkin’s lymphoma (NHL), ALL, AML, CLL, chronic myeloid leukemia (CML), blastic plasmacytoid dendritic cell neoplasm (BPDCN), and multiple myeloma (MM). HemaCisDB is a user-friendly and interactive database allowing users to query, browse, and visualize cis-regulatory regions in datasets of interest. It provides comprehensive quality control (QC) metrics and analytical modules including TF footprinting, SE identification/ranking, and CRC analysis. HemaCisDB also allows users to compare TF footprints among user-defined datasets to explore TF binding dynamics across different disease types or conditions. By incorporating the latest epigenomic profiling datasets into HemaCisDB, we provide a comprehensive resource to facilitate the study of CREs and their functions in hematopoietic malignancies.

### Database construction

#### Data collection

In HemaCisDB, we manually curated 1643 publicly available ATAC-seq (n = 922), DNase-seq (n = 190), and H3K27ac ChIP-seq (n = 531) datasets from blood cancer cell lines and patient samples with hematopoietic malignancies. ATAC-seq/DNase-seq or H3K27ac ChIP-seq datasets were explored using the following keywords: “ATAC-seq” (or “DNase-seq” or “H3K27ac”), “*Homo sapiens*”, “AML” (or “acute myeloid leukemia”), “CML” (or “chronic myeloid leukemia”), “ALL” (or “acute lymphocytic leukemia”), “CLL” (or “chronic lymphocytic leukemia”), “MM” (or “multiple myeloma”) and “Lymphoma” against the Gene Expression Omnibus (GEO) database (http://www.ncbi.nlm.nih.gov/geo). All datasets were manually reviewed and irrelevant data, such as single-cell ATAC-seq, were removed from the final list.

#### ATAC-seq/DNase-seq and H3K27ac ChIP-seq data processing

Raw reads were trimmed by TrimGalore (version 0.6.7) to remove adaptors and low-quality sequences. Trimmed reads were then mapped to human genome assembly (GRCh38 GENCODE release 39) using bowtie2 (version 2.2.5) with default parameters [25]. For ATAC-seq/DNase-seq data, fragments mapped to mitochondria DNA or with length greater than 2000bp or less than 38bp were filtered out by samtools (version 1.6) [26]. Similarly, only fragments less than 2kb in H3K27ac ChIP-seq data were kept for further analysis. Picard (version 2.27.5) was applied to remove duplicated reads. Accessible chromatin regions or regions with H3K27ac modification were identified by MACS2 (version 2.2.1.4) with parameters of ‘--nomodel --shift −100 --extsize 200’ and default settings respectively [27]. ATAC-seq/DNase-seq and H3K27ac ChIP-seq peaks overlapped with the curated blacklisted regions (https://github.com/Boyle-Lab/Blacklist/tree/master/lists) were excluded from our database [28].

#### ATAC-seq/DNase-seq quality control

R packages ATACseqQC [29] and encodeChIPqc were used to evaluate ATAC-seq/DNase-seq data quality. Five QC metrics, including sequencing depth and library complexity assessment, size distribution of library insert fragments, transcription start site (TSS) enrichment score, nucleosome density distribution around TSS and Fraction of Reads in Peaks (FRiP) were used in HemaCisDB. High-quality ATAC-seq data should 1) have large proportion of insert fragments less than 100bp which represents nucleosome-free region, and fragment size distribution exhibits a clear periodicity corresponding to nucleosome-free regions and the occupancy of mono-nucleosomes, di-nucleosomes, and tri-nucleosomes; 2) show good TSS enrichment; 3) display strong enrichment of nucleosome-free fragments at TSS and nucleosome-bound fragments upstream and downstream of the active TSSs; 4) show high FRiP score with high fraction of mapped reads falling into the peak regions.

#### H3K27ac ChIP-seq quality control

H3K27ac ChIP-seq data quality was assessed using R package ChIPQC [30]. FRiP, relative strand cross-correlation coefficient (RelCC), fragment length, and standardized standard deviation (SSD) were included in the QC metrics. FRiP, as calculated in ATAC-seq, represents mapped reads within called peaks, and high FRiP indicates high signal-to-noise ratio of the investigated data. RelCC is another “signal-to-noise” measurement which is determined from the calculation of strand cross-correlation, and a RelCC value greater than 1 suggests good signal-to-noise ratio. Fragment length estimated from ChIP-seq data should be consistent with fragment size selected in library preparation step. SSD, calculated from the standard deviation of signal pile-up across chromosomes normalized to the total number of reads sequenced, represents the uniformity of coverage of reads along the genome with a higher SSD indicative of better enrichment.

#### Annotation and functional enrichment of H3K27ac ChIP-seq and ATAC-seq/DNase-seq peaks

R packages ChIPseeker was used to annotate ChIP-seq and ATAC-seq/DNase-seq peaks [31]. The nearest genes of the peaks, the distances between the peaks to the TSSs of their nearest genes, and the genomic regions overlapped with the peaks were reported. The priorities of genomic annotation are as follows: promoter, 5’ UTR, 3’ UTR, exon, intron, downstream, and intergenic. Functional enrichment analysis was performed by R package clusterProfiler [32] based on the annotated nearest genes. Enriched pathways in biological processes of Gene Ontology (GO) and Kyoto Encyclopedia of Genes and Genomes (KEGG) were identified.

#### TF footprint analysis

In ATAC-seq/DNase-seq, DNA bases bound by TFs are protected from Tn5/DNase I cleavage within the binding site, resulting in a relative depletion of signal within the open chromatin region, known as footprints. TF footprinting analysis allows for the prediction of precise TF binding sites. TOBIAS, which incorporates bias correction and footprinting, was used to perform TF footprinting analysis in HemaCisDB [33]. For each motif obtained from JASPAR, the number of binding sites found in peak regions and mean footprint score for all TF binding sites were reported. A higher mean score indicates a clearer footprint of a TF. An aggregate plot across all sites for each TF was also generated for footprint visualization. Comparison of TF footprints across different samples and different conditions was also performed by TOBIAS, which allows users to investigate differential binding of TFs between different user-defined samples or conditions.

#### Super-enhancer Identification

H3K27ac peaks were used to define SE boundaries, with the following filtering criteria: (1) excluding H3K27ac peaks that overlapped with ENCODE blacklisted genomic regions [28], and (2) excluding H3K27ac peaks that were located within ± 2 kb region of any RefSeq annotated gene promoter. SEs were then identified based on the H3K27ac ChIP-seq signals using the ROSE algorithm2 with the default parameters [34].

#### CRC analysis

CRC analysis was performed using Python package coltron on samples with paired H3K27ac ChIP-seq and ATAC-seq data [35]. The command line to run coltron is as follows: coltron -e [output Enhancer file from ROSE] -b [H3K27ac ChIP-seq bam file] -g “37” -n [Name] -s [Subpeak file] –promoter = True -o [Output] –promoter = True. And subpeaks in the above command were defined as ATAC-seq peaks that are overlapped with H3K27ac enhancer regions.

Specifically, coltron applies the following strategy to identify autoregulatory and interconnected TF networks. The top 1000 enhancers ranked by H3K27ac signal were assigned to nearby genes whose promoters are highly acetylated. The underlying sequences of ATAC-seq peaks within H3K27ac enhancer regions were extracted and FIMO (version 4.91) was employed to search TF node binding sites with position-weight matrices from Transfac and Jaspar. An edge will be drawn between TF A and TF B if the motif of TF A is identified in an enhancer assigned to regulate TF B. In networks, the in-degree of TF A is defined as the number of TF nodes within any enhancers that regulate TF A. And the out-degree of TF A is described as the number of TF nodes regulated by an enhancer with the motif of TF A.

#### SNPs and eQTL annotation

Common single nucleotide polymorphisms (SNPs) and risk SNPs within accessible chromatin regions or enhancer regions were annotated. Common SNPs were obtained from dbSNP 156 [36], and SNPs with minimum allele frequency (MAF) < 0.01 were filtered out. Risk SNPs were retrieved from the GWAS Catalog [37], and risk SNPs associated with blood disorders were manually curated from all risk SNPs. Pairwise linkage disequilibrium (LD) was computed among SNPs with MAF greater than 0.05. PLINK (version 2.0) [38] was employed to calculate the LD SNPs (r2 = 0.8) in five super-populations, including African (AFR), Ad Mixed American (AMR), East Asian (EAS), European (EUR) and South Asian (SAS). Expression quantitative trait loci (eQTL) and eQTL-gene pairs were acquired from PancanQTL [39] and GTEx (version 8.0) [40].

#### Web portal

VueJS (version 2.6.10) (https://vuejs.org) and Element UI (version 2.13.2) (https://element.eleme.io/) were used to develop the front end of server, and the back end was built in Python using the Django web framework (version 3.2.8). The server is hosted on a Linux Ubuntu 20.04 server running Nginx/1.18.0. Files of the raw data matrix, pre-processed data and intermediate results were stored in the local Ext4 file system, and MySQL (version 8.0.31) database was used to store metadata and analysis results.

### Database content and usage

#### Scheme of HemaCisDB

HemaCisDB currently contains 922 ATAC-seq, 190 DNase-seq, and 531 H3K27ac ChIP-seq datasets from 133 publications (**Figure 1**). We performed QC, peak identification, and peak annotation for each ATAC-seq, DNase-seq, and H3K27ac ChIP-seq dataset (Figure 1). As TF-bound DNA sequences are protected from Tn5-mediated transposition or DNase I digestion, leaving behind “footprints”, ATAC-seq and DNase-seq were proposed to be useful tools to infer genome-wide TF-bound locations [33, 41]. HemaCisDB provides footprinting analysis for each ATAC-seq/DNase-seq dataset and allows users to explore TF binding information across diverse subtypes of hematopoietic malignancies (Figure 1).

**Figure 1.**
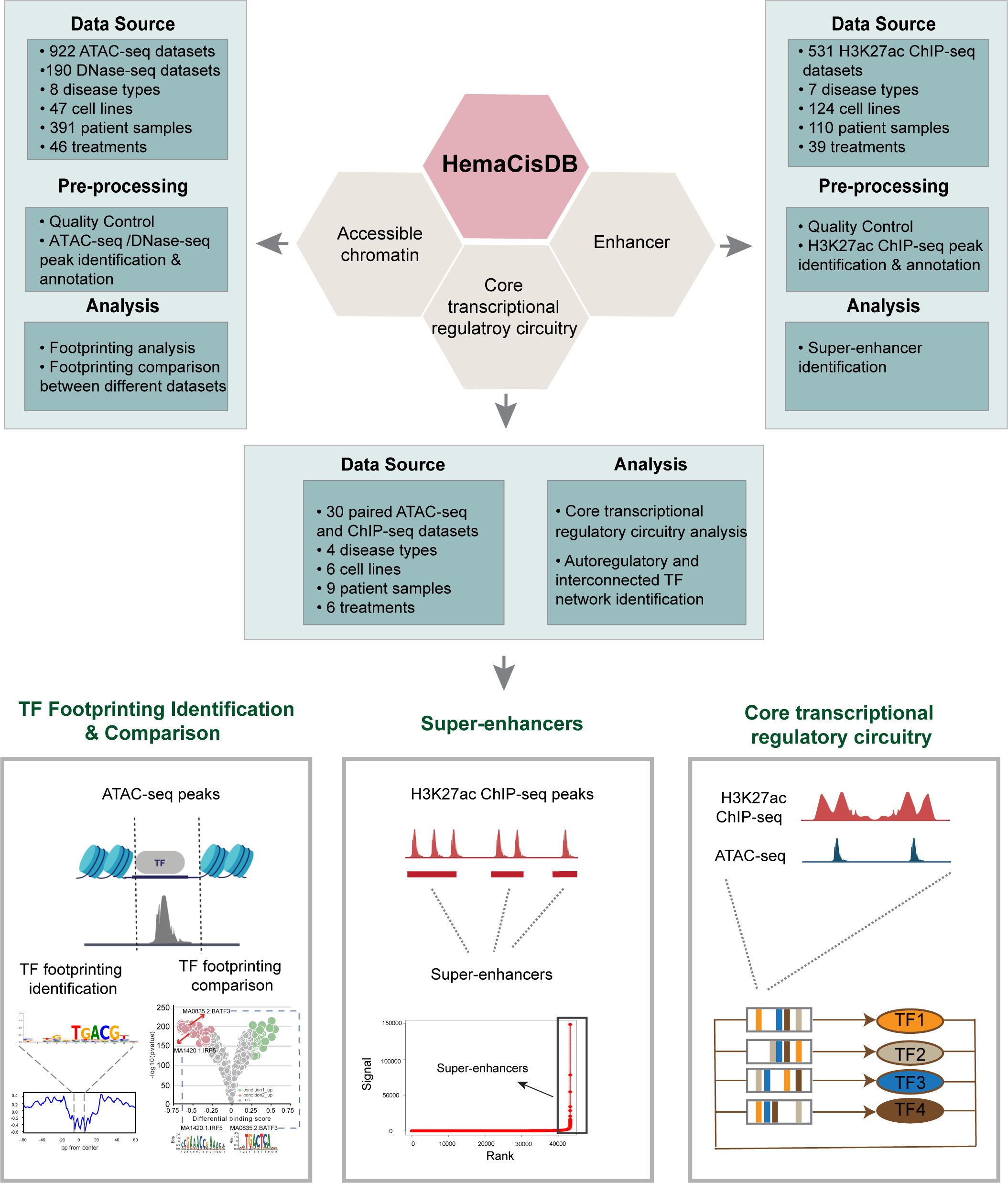
Scheme of HemaCisDB. HemaCisDB contains three main modules: accessible chromatin, enhancer, and core transcriptional regulatory circuitry. ATAC-seq/DNase-seq data was employed to identify open chromatin regions, and H3K27ac ChIP-seq data was utilized for enhancer identification. The core transcriptional regulatory circuitry was identified from paired ATAC-seq and H3K27ac ChIP-seq data. Quality control, peak identification, and peak annotation were performed for each ATAC-seq/DNase-seq and H3K27ac ChIP-seq dataset. HemaCisDB provides analytical functions within these three modules. TF footprinting and comparison are available in the Accessible chromatin module, super-enhancer identification is provided in the Enhancer module, and autoregulatory and interconnected TF networks can be identified in the Core transcriptional regulatory circuitry module. ATAC-seq, the assay of transposase accessible chromatin with sequencing; DNase-Seq, DNase I hypersensitive sites sequencing; H3K27ac, H3K27 acetylation; ChIP-seq, chromatin immunoprecipitation followed by sequencing; TF, transcription factor.

A SE, which comprises clusters of enhancer elements, is another important genomic feature that plays critical roles in driving cell-type-specific and disease-associated gene expression [42]. To uncover SEs that control gene programs related to different blood disorders, HemaCisDB enables SE identification by stitching closely distributed enhancers determined from H3K27ac ChIP-seq (Figure 1). For samples with paired ATAC-seq and H3K27ac ChIP-seq data, CRC analysis was performed to identify autoregulatory and interconnected TFs that regulate lineage-specific and disease-associated genes (Figure 1), thus providing information for elucidating malignant characteristics of different blood cancer subtypes [43, 44].

#### Datasets in HemaCisDB

HemaCisDB contains ATAC-seq, DNase-seq, and H3K27ac ChIP-seq datasets from 294 healthy donor samples and 1349 disease-related samples (**Figure 2A**). Disease-related datasets include 8 subtypes of blood disorders across myeloid and lymphoid neoplasms, including AML, CML, ALL, CLL, MM, NHL, HL, and BPDCN (Figure 2A). NHL collected in the database consists of Burkitt’s lymphoma (BL), DLBCL and Mantle Cell Lymphoma (MCL). The datasets collected in HemaCisDB are from cell lines or patient samples. 43, 19, 53, and 14 cell lines were collected from diseases of myeloid leukemia, lymphocytic leukemia, lymphoma, and MM respectively (Figure 2B, Figure S1A). Patient samples are mainly derived from bone marrow or peripheral blood (Figure 2B). Treatment information of each dataset is also available in HemaCisDB, allowing users to investigate epigenetic landscape change under different treatment conditions. Treatment categories mainly include genetic or pharmacological perturbations targeting epigenetic factors, TFs, and signaling pathways (Figure 2C, Figure S1B).

**Figure 2.**
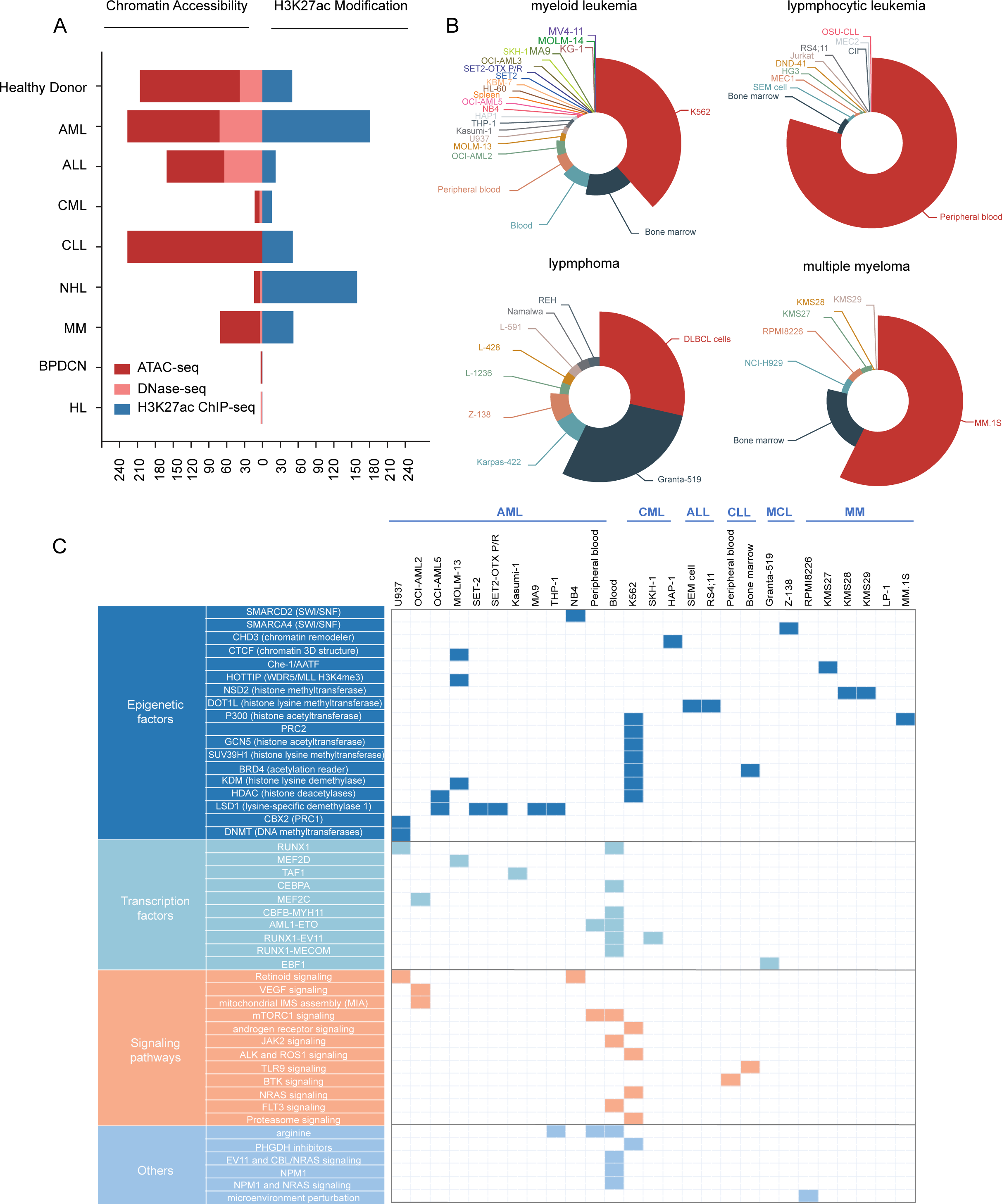
Statistics of datasets in HemaCisDB. **A**. Number of ATAC-seq, DNase-seq, and H3K27ac ChIP-seq datasets summarized by disease type. **B.** Number of ATAC-seq/DNase-seq datasets summarized by sample source. **C.** Summarization of genetic or drug perturbations across different sample sources for ATAC-seq/DNase-seq. AML, acute myeloid leukemia; ALL, acute lymphocytic leukemia; CML, chronic myeloid leukemia; CLL, chronic lymphocytic leukemia; NHL, non-Hodgkin lymphoma; MM, multiple myeloma; BPDCN, blastic plasmacytoid dendritic cell neoplasm; HL, Hodgkin lymphoma; MCL, mantle cell lymphoma.

#### A user-friendly interface for browsing, querying, and downloading datasets

HemaCisDB allows users to browse samples and filter datasets by disease type, biosample type, and biosample source (**Figure 3A**, Figure S2A). A search option is also supported, allowing users to query a dataset through corresponding study ID, sample ID, disease type, cell line name, or other related information (Figure 3A, Figure S2A). Users can visualize detailed information for each ATAC-seq or H3K27ac ChIP-seq dataset, including QC metrics, peak coordinates, and peak annotations (Figure 3B, Figure S2B). All peak regions and peak information are available for download from HemaCisDB (Figure 3C, Figure S2C). Furthermore, peak regions derived from ATAC-seq or H3K27ac ChIP-seq across different datasets have been integrated and annotated. Detailed information for each region is accessible through the ‘Accessible regions’ or ‘Enhancer regions’ tabs under ‘Accessible chromatin’ or ‘Enhancer’ respectively. Users can explore comprehensive annotations for each region, including common SNPs, risk SNPs, SNPs associated with blood disorders, eQTLs, and LD SNPs, by clicking on the ‘Region ID’ (Figure 3D and Figure S2D). In addition, each region is annotated to indicate whether it is shared among multiple disease types or exclusive to a specific disease type.

**Figure 3.**
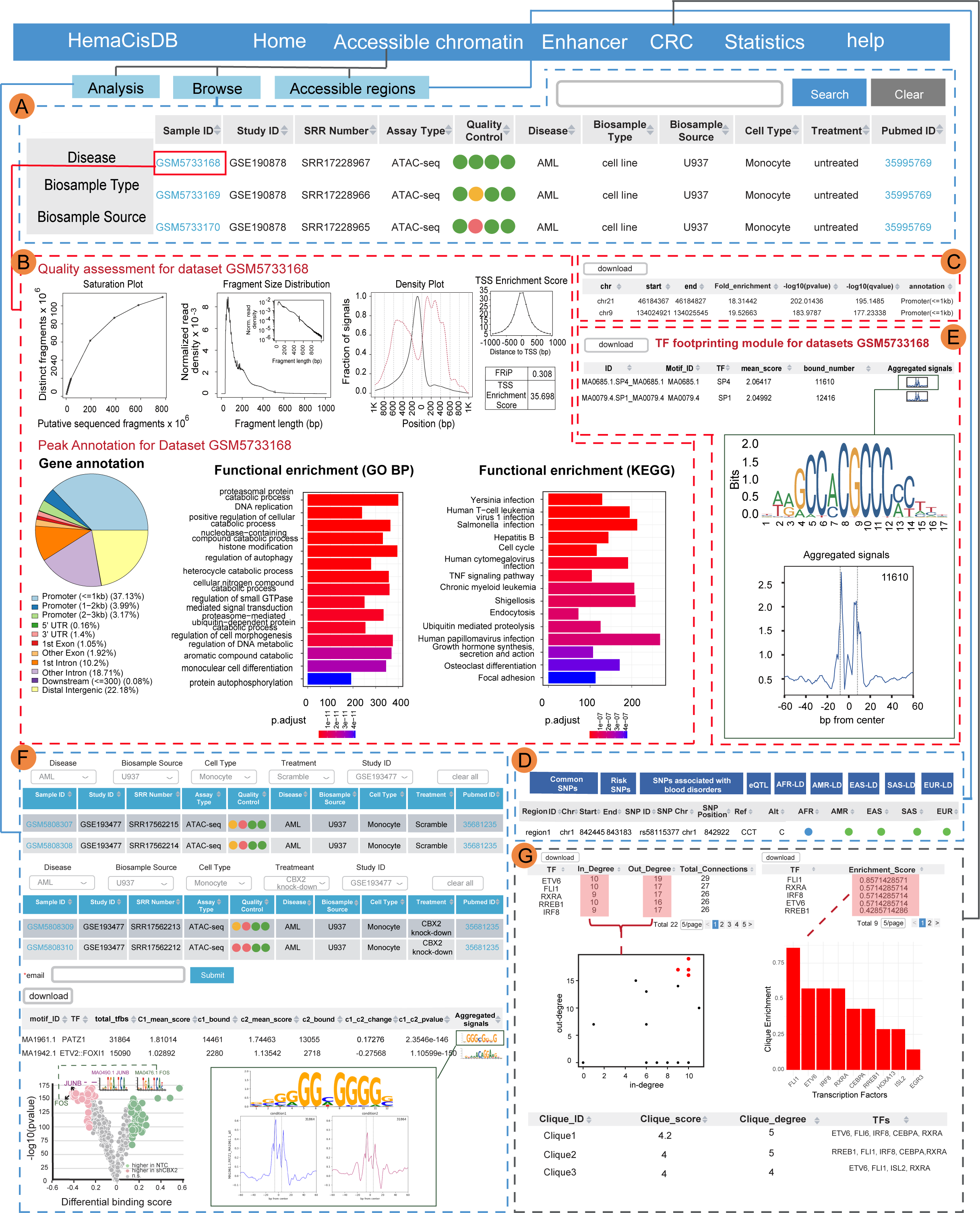
Data browsing and application modules for ATAC-seq/DNase-seq data. **A.** Data browsing page for ATAC-seq/DNase-seq data. **B.** Quality assessment and functional annotation of peaks for selected ATAC-seq/DNase-seq dataset. **C.** ATAC-seq/DNase-seq peaks identified from selected dataset. Coordinates, fold enrichment, *P* value, *Q* value, and annotation of each peak are reported. **D.** Common SNPs, risk SNPs, and risk SNPs associated with blood disorders that overlap each accessible region are reported. For each common SNP, corresponding eQTL and LD SNPs in five super-populations (AFR, AMR, EAS, EUR, and SAS) are also reported. **E.** TF footprint inference page. The information of motif ID, TF name, mean footprint score across all TF binding sites, and the number of predicted binding sites in peak regions is reported. The aggregation plot and motif logo for each TF can be visualized. **F.** TF footprint comparison page. Disease type, biosample source, cell type, treatment, and study ID can be used to filter and select datasets to be compared. The link of comparison results will be sent to users by email. Differential binding of each TF between different samples or different conditions is reported. Each point in the scatter plot represents a TF, with x axis as the binding differences of a TF between two different user-selected samples and y axis as -log10 (*P* value). **G.** CRC analysis page. The information of TF in-degrees, TF out-degrees, TF enrichment scores and top scoring TF cliques is reported. Each point in the scatter plot represents a TF, with x axis as the in-degree of a TF and y axis as the out-degree of a TF. Bar plot shows TFs ranked by clique enrichment scores. SNP, single nucleotide polymorphism; eQTL, expression quantitative trait loci; LD, linkage disequilibrium; AFR, African; AMR, Ad Mixed American; EAS, East Asian; EUR, European; SAS, South Asian; CRC, core transcriptional regulatory circuitry.

#### Application modules

Four analytic modules were developed in HemaCisDB.

##### TF footprinting module

HemaCisDB provides TF footprint inference within accessible chromatin regions (Figure 3E). For each TF, the number of predicted binding sites in peak regions and mean footprint score across all TF binding sites are reported. Users can visualize footprints and motif logo for each TF via aggregation plot and sequence logo plot.

##### TF footprinting comparison module

HemaCisDB also allows users to compare TF footprints across different samples (Figure 3F). HemaCisDB facilitates the comparative analysis of TF footprints between different samples (Figure 3F). Users can select from the drop-down menu of disease types, cell types, or treatment conditions to identify samples for comparisons. Output with differentially bound TFs between different samples will be generated. Differences and *P* values of each TF are reported, and users can visualize the summarized results through a volcano plot. For each TF, HemaCisDB also provides aggregate plots across different regions between different samples.

##### Super-enhancer module

This module shows detailed SE information identified from H3K27ac ChIP-seq datasets (Figure S2E). The coordinates, constitute size, signal rank, and annotation of each SE are reported. For each dataset, HemaCisDB also generates the plot of stitched enhancers ranked by H3K27ac ChIP signals, allowing users to visualize the distribution of SEs.

##### CRC analysis module

HemaCisDB provides CRC analysis to identify autoregulatory and interconnected TF networks for each sample with paired H3K27ac ChIP-seq and ATAC-seq data (Figure 3G). This module shows TF degrees, TF networks, clique enrichment of TFs, and detailed information of cliques with top clique scores. The degree for a given SE-regulated TF is defined as the sum of in-degree and out-degree of the TF. In-degree is calculated as the number of SE-regulated TFs bound to the assigned SEs of the given TF. And out-degree is described as the number of TF-associated SEs bound by the given SE-regulated TF. A clique in TF networks is defined as a subnetwork containing at least four TF nodes that are connected to themselves and all other TFs within the network. ‘Clique enrichment’ of a given TF is defined as the percentage of cliques with that TF, and ‘clique score’ is calculated as the average of clique enrichment. In-degree and out-degree of each TF are reported and summarized by scatter plots (Figure 3G). Clique enrichment of TFs is listed and TFs with top clique enrichment scores are shown in ordered bar plots (Figure 3G). Users can visualize detailed information of top scoring cliques, including clique scores, clique degrees, and all TFs in that clique.

#### CRE landscape across different subtypes of hematopoietic malignancies

Integrative analysis was conducted to categorize accessible chromatin/enhancer regions, obtained from ATAC-seq/DNase-seq and H3K27ac ChIP-seq data across all compiled datasets, into common regions shared by multiple disease types and disease-specific regions (**Figure 4A–B**). Hierarchical clustering of the ATAC-seq peak/DNase-seq and H3K27ac ChIP-seq peak regions showed that cell lines and primary samples from myeloid neoplasms were grouped together, and samples derived from lymphoid malignancies exhibited higher similarities (Figure 4C–D). Thus, CRE landscape can successfully capture the information of lineages and disease types.

**Figure 4.**
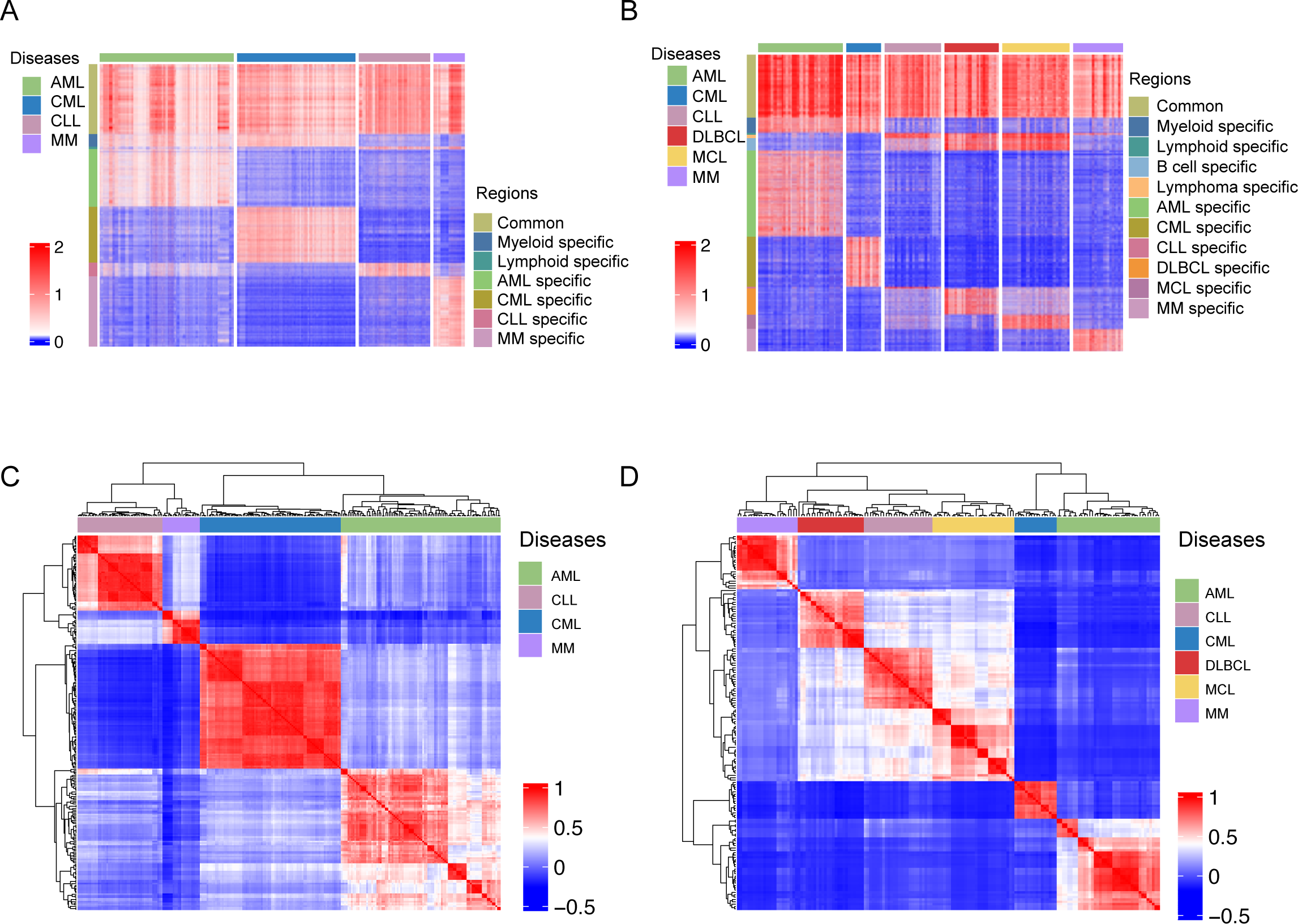
ATAC-seq/DNase-seq and H3K27ac ChIP-seq profiles across different types of hematopoietic malignancies. **A.** ATAC-seq/DNase-seq signals within common and disease-specific peaks across different types of blood cancers. **B.** H3K27ac ChIP-seq signals within common and disease-specific peaks across different types of blood cancers. **C.** Clustering analysis of ATAC-seq profiles across different cell lines, cell types, or disease lineages. **D.** Clustering analysis of H3K27ac ChIP-seq profiles across different cell lines, cell types, or disease lineages. DLBCL, diffuse large B cell lymphoma.

#### Demonstrations of HemaCisDB usage

CREs studies in cancer can reveal critical transcriptional regulators and uncover the underlying molecular mechanisms driving cancer pathogenesis. Here, we showcase how the HemaCisDB database can be utilized to achieve these aims. Previous studies showed that CREs linked to lineage-specific genes are commonly dysregulated in hematopoietic malignancies [9, 15]. HemaCisDB is a useful resource that facilitates the identification of cell-type-specific CREs and prediction of prospective regulators in user-defined diseases. CLL is a type of adult leukemia that originates primarily from malignant B cells. Analysis using HemaCisDB revealed that SEs associated with B-cell-specific TFs *PAX5* and *BCL2* are exclusively present in CLL compared to AML, as evidenced by enriched H3K27ac ChIP-seq and ATAC-seq signals in CLL cell line MEC1, and depleted signals in AML cell line MOLM-13 (**Figure 5A**). Conversely, SEs associated with TFs involved in myeloid development, such as *RUNX1* and *CEBPA*, exhibited enriched H3K27ac ChIP-seq and ATAC-seq signals solely in AML (Figure 5A). Furthermore, CRC analysis in MEC1 highlighted PAX5 as a crucial transcriptional regulator with high TF degrees, high clique enrichment score, and involvement in top scoring auto-regulatory TF networks (Figure 5B–D), signifying its potential roles in CLL pathogenesis. These findings align with a previously published study indicating that PAX5 serves as a core regulator in CLL regulatory networks and plays an indispensable role in the maintenance of CLL cell viability [45].

**Figure 5.**
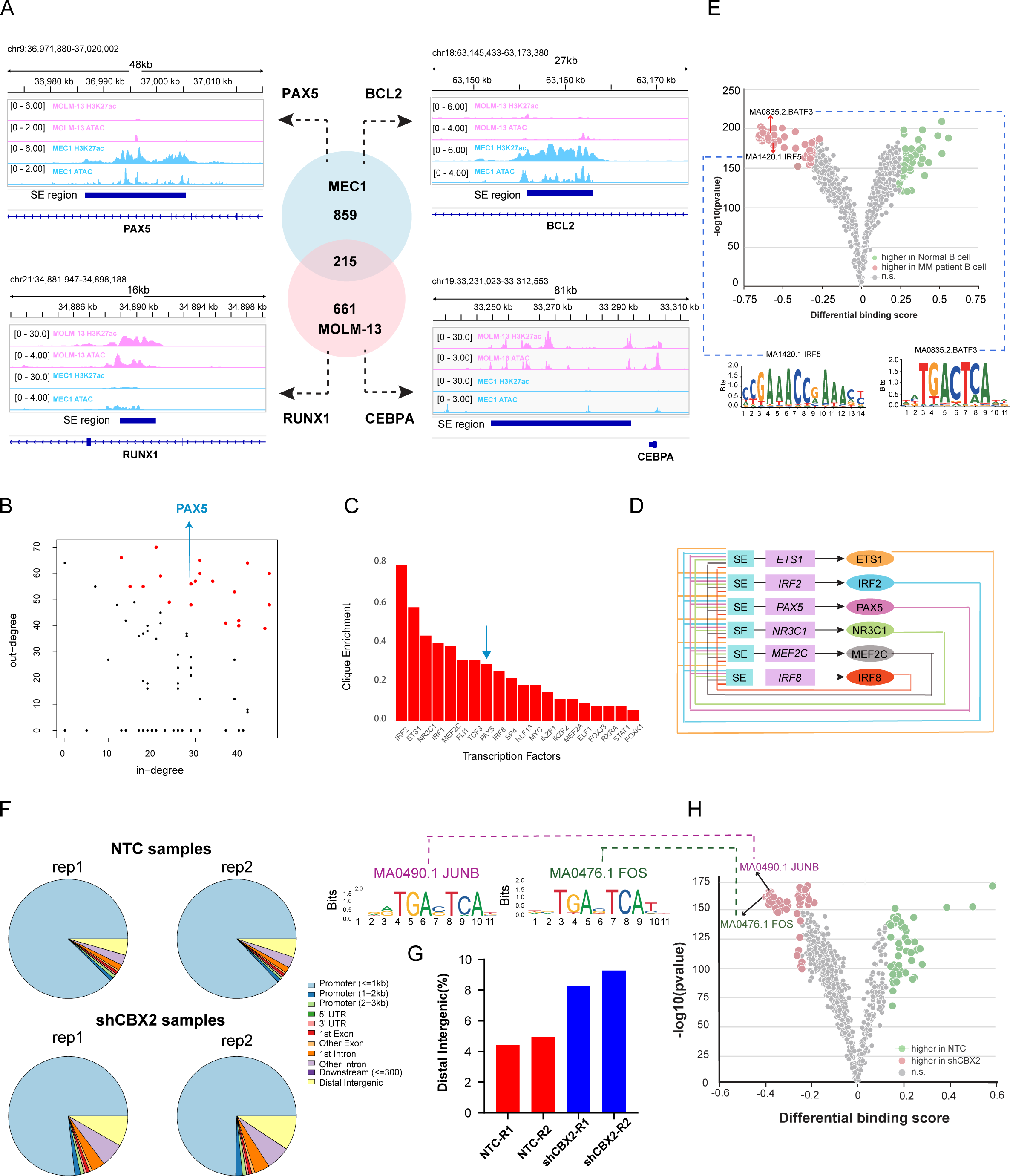
Demonstration of HemaCisDB usage by case studies. **A.** SE identification in CLL cell line MEC1 and AML cell line MOLM-13. SEs were identified using the analytic module of *Super-enhancer Identification* in HemaCisDB. **B.** TF degree analysis performed by *CRC analysis module* of HemaCisDB highlights PAX5 as a critical TF in CLL cell line MEC1 with relatively high TF in-degrees and TF out-degrees. Each point in the scatter plot represents a TF, with x axis as the in-degree of a TF and y axis as the out-degree of a TF. **C.** TF enrichment score calculated by *CRC analysis module* of HemaCisDB highlights PAX5 as an important TF in CLL cell line MEC1 with relatively high TF enrichment score. Bar plot shows TFs ranked by clique enrichment scores. **D.** PAX5 is involved in top scoring TF clique in CLL cell line MEC1. TF cliques were identified from *CRC analysis module* of HemaCisDB. E. Analysis performed by *TF footprinting comparison* module of HemaCisDB shows significant enrichments of IRF5 and BATF3 motifs at open chromatin regions in MM tumor cells compared to normal B cells. Each point in the scatter plot represents a TF motif collected from JASPAR. **F.** Annotations of ATAC-seq peaks for AML cell lines transduced with shCBX2 or non-targeting controls. NTC denotes the non-targeting controls, and peak annotation for each ATAC-seq dataset is available at HemaCisDB. **G**. Percentage of ATAC-seq peaks located at the distal intergenic regions in AML cell lines transduced with shCBX2 or non-targeting controls. **H.** Analysis performed by *TF footprinting comparison module* of HemaCisDB shows significant enrichments of motifs associated with TFs JUNB and FOS at open chromatin sites in cell lines with CBX2 silencing. Each point in the scatter plot represents a TF motif collected from JASPAR. SE, super-enhancer.

Comparing malignant cells to their normal counterparts from healthy donors can offer valuable insights into the molecular mechanisms underlying cancer pathogenesis. HemaCisDB has curated ATAC/DNase-seq and H3K27ac ChIP-seq data from various purified cell types of healthy donors. Additionally, the TF footprinting comparison module in HemaCisDB allows for the comparison of TF footprints across different samples. We demonstrated the potential of HemaCisDB in identifying TF candidates that drive cancer pathogenesis by comparing TF footprints using ATAC-seq data of fluorescence-activated cell sorting (FACS)-purified myeloma cells from multiple myeloma patients and normal B cells from primary human blood cells. This analysis revealed a significant enrichment of the IRF5 motif within open chromatin regions in MM cells (Figure 5E), consistent with previous findings that identified IRF5 regulon as a cancer-specific regulon with higher activity in MM cells versus normal B cells using scRNA-seq and scATAC-seq [46]. Additionally, the BATF3 motif was also found to be enriched within open chromatin regions in MM patient tumor cells (Figure 5E). The patient sample used for comparison harbors the t(11;14) translocation, aligning with prior findings that BATF3 expression co-occurs with t(11;14) translocation [47]. BATF3 expression has also been reported to be associated with worse progression-free survival (PFS), overall survival, and increasing clinical stages in MM [47]. These imply that IRF5 and BATF3, identified through HemaCisDB footprinting comparison, could potentially be critical TFs driving MM pathogenesis.

The employment of genetic perturbation methods, such as CRISPR knockout and short hairpin RNAs (shRNAs), has facilitated gene function investigations. HemaCisDB has also curated treatment information from collected datasets, including genetic or drug perturbations of genes that encode epigenetic factors, TFs, or proteins in signaling pathways. This provides a valuable resource to study how dysfunction of different genes contributes to the pathogenesis of hematopoietic malignancies. Here, we demonstrate the utility of HemaCisDB in investigating how gene perturbations contribute to disease development through CRE deregulations. CBX family of proteins are canonical PRC1 subunits, which can read and bind to H3K27me3 modifications [48]. This, in turn, facilitates the recruitment of canonical PRC1 to PRC2 target genes, thereby leading to chromatin condensation and transcriptional repression. CBX2 overexpression has been implicated in promoting the proliferation of cancer cells [49]. HemaCisDB revealed that CBX2 silencing leads to an increase in open chromatin regions at distal intergenic regions in the AML cell line U937 (Figure 5F–G). Furthermore, the footprinting comparison of AML cell lines transduced with shCBX2 and non-targeting controls revealed that motifs for AP-1 TFs, such as JUNB and FOS, were significantly enriched in open chromatin sites in cell lines with CBX2 silencing (Figure 5H). AP-1 TFs, which act as effectors of receptor tyrosine kinase signaling and growth factor, are activated by the Ras/MAPK pathway [50], suggesting that CBX2 might promote AML through MAPK-associated regulatory sites. These results are consistent with a previous study that demonstrated that silencing CBX2 results in epigenetic reprogramming of regulatory sites associated with p38 MAPK, leading to gene expression deregulation [51].

Collectively, HemaCisDB offers an interactive platform for CRE characterization, facilitating mechanistic investigations of transcriptional regulation in hematopoietic malignancies.

## Discussion

CRE landscape can serve as a useful biomarker to identify blood cancer subtypes. Dysregulation of CREs often contributes to pathogenesis and drug resistance of hematopoietic malignancies [52]. The comprehensive and unbiased characterization of CREs facilitate the analysis of putative cell-of-origin and/or targetable vulnerabilities of blood disorders. Databases, such as Cistrome, ATACdb, dbSUPER, SEA, SEdb, and Cancer CRC, have been developed to characterize CREs across various cell types and tissues in different types of cancers. However, these resources were mainly designed for pan-cancer analysis and only encompass a limited number of datasets relevant to hematopoietic malignancies (Table S1). Thus, we developed HemaCisDB, a comprehensive and interactive data portal with specific focus on CREs in hematopoietic malignancies. HemaCisDB supports disease-specific queries in blood cancers. In contrast to many previously published databases that solely collected ATAC-seq/DNase-seq or H3K27ac ChIP-seq data, HemaCisDB provides the most extensive collection of datasets and CRE profiles related to blood cancers (Table S1). This is achieved by integrating both ATAC-seq/DNase-seq and H3K27ac data, enabling concurrent characterization of open chromatin regions, enhancers/SEs, and CRCs (Table S1). HemaCisDB provides comprehensive QC metrics, facilitating the QC assessment for each dataset. Furthermore, HemaciDB profiled treatment conditions, allowing investigations into how different treatments affect transcriptional regulation in blood disorders (Table S1). Additionally, HemaCisDB provides detailed annotations for each open chromatin and enhancer region, including genomic features, SNPs, eQTLs, and an indication of whether a region is shared among multiple disease types or exclusive to a specific disease type (Table S1). These comprehensive annotations allow for in-depth functional exploration of each region. Taken together, HemaCisDB serves as a resource facilitating the hematology community in characterizing CREs and their roles in disease pathophysiology.

## Data availability

HemaCisDB is accessible at https://hemacisdb.chinablood.com.cn/.

## CRediT author statement

**Xinping Cai**: Methodology, Software, Investigation, Data curation, Writing - original draft, Writing - review & editing. **Qianru Zhang**: Investigation, Data curation. **Bolin Liu**: Data curation. **Lu Sun**: Data curation. **Yuxuan Liu**: Conceptualization, Methodology, Writing - original draft, Writing - review & editing, Supervision, Project administration, Funding acquisition. All authors read and approved the final manuscript.

## Competing interests

The authors have declared no competing interests.

## Acknowledgements

We are grateful to Jian Xu at St. Jude Children’s Research Hospital for helpful discussions and proofreading the manuscript. This work was supported by the Non-profit Central Research Institute Fund of Chinese Academy of Medical Sciences (Grant No. 2022-RC310-03), Chinese Academy of Medical Sciences Innovation Fund for Medical Sciences (CIFMS) (Grant No. 2022-I2M-1-022), State Key Laboratory of Experimental Hematology Research Grant (Grant No. Z22-01), and Biomedical High Performance Computing Platform, Chinese Academy of Medical Sciences. We also thank National SuperComputer Center in Tianjin for their help in developing the web interface for the HemaCisDB.

## Supplementary material

**Supplementary Figure S1 Statistics of H3K27ac ChIP-seq datasets in HemaCisDB A.** Number of H3K27ac ChIP-seq datasets summarized by sample source. **B.** Summarization of genetic or drug perturbations across different sample sources for H3K27ac ChIP-seq. H3K27ac, H3K27 acetylation; ChIP-seq, chromatin immunoprecipitation followed by sequencing.

**Supplementary Figure S2 Data browsing and application modules for H3K27ac ChIP-seq data A.** Data browsing page for H3K27ac ChIP-seq data. **B.** Quality assessment and functional annotation of peaks for selected H3K27ac ChIP-seq dataset. **C.** H3K27ac ChIP-seq peaks identified from selected dataset. Coordinates, fold enrichment, *P* value, *Q* value, and annotation of each peak are reported. **D.** Common SNPs, risk SNPs, and risk SNPs associated with blood disorders that overlap each enhancer region are reported. For each common SNP, corresponding eQTL and LD SNPs in five super-populations (AFR, AMR, EAS, EUR, and SAS) are also reported. **E.** Super-enhancer identification page. Coordinates, the number of enhancers stitched together, constituent size, signal, rank, overlap and proximal genes of each SE are reported. SE distribution was plotted with x-axis as the rank of SEs and y-axis as the level of signals of SEs. SNP, single nucleotide polymorphism; eQTL, expression quantitative trait loci; LD, linkage disequilibrium; AFR, African; AMR, Ad Mixed American; EAS, East Asian; EUR, European; SAS, South Asian. SE, super-enhancer.

**Table S1 A comprehensive comparison between HemaCisDB and other databases**

