## Supplementary figures and images for "HemaCisDB: An Interactive Database for Analyzing Cis-Regulatory Elements Across Hematopoietic Malignancies"

### Supplemental Figure 1

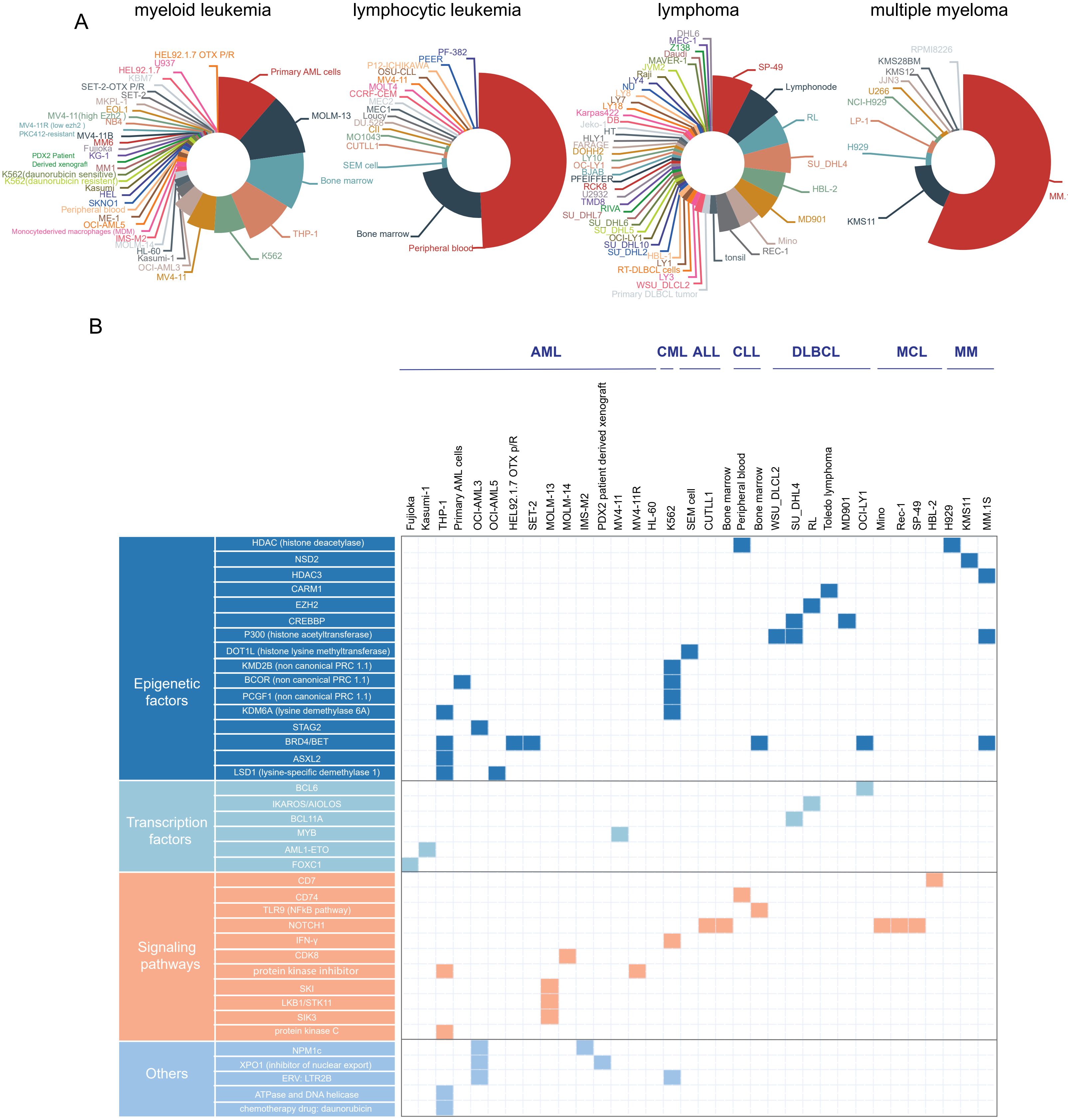

### Supplemental Figure 2

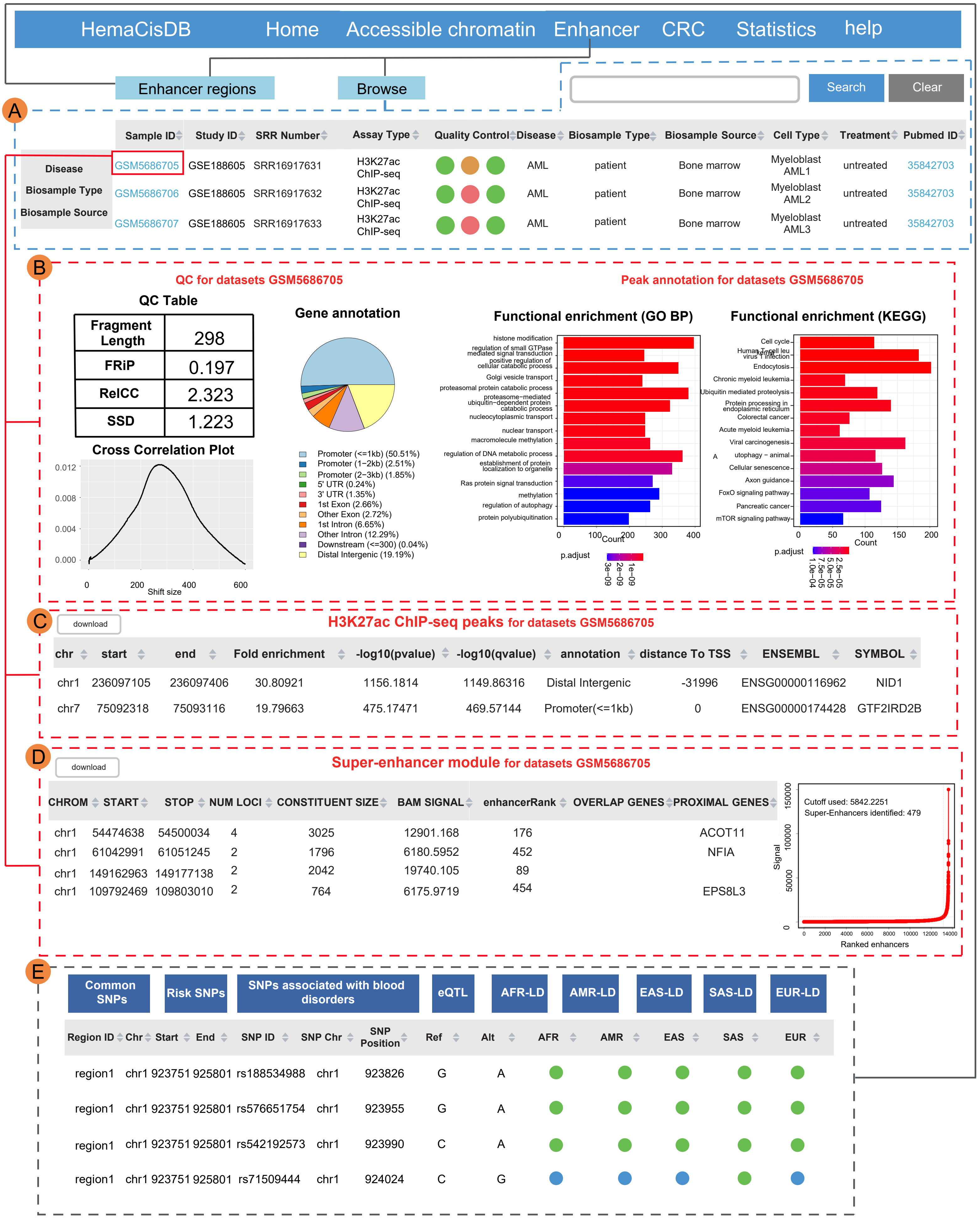
