## Supplemental Table 1 for "HemaCisDB: An Interactive Database for Analyzing Cis-Regulatory Elements Across Hematopoietic Malignancies"

| **Function type** | **Data type/Specific function** | **HemaCis**  **DB** | **Cistrome**  **DB(v.3.0)** | **ATAC**  **db** | **db**  **SUPER** | **SEA**  **(v.3.0)** | **SEdb**  **(v.2.0)** | **Cancer CRC** |
| --- | --- | --- | --- | --- | --- | --- | --- | --- |
| **Data Source** | Number of blood cancer types | 10 | 8 | 8 | 4 | 6 | 8 | 0 |
|  | Number of ATAC-seq/DNase-seq samples^a^ | 856 | 502 | 137 | 0 | 0 | 0 | 0 |
|  | Number of H3K27ac ChIP-seq samples^a^ | 481 | 316 | 0 | 12 | 9 | 192 | 0 |
| **Data browse** | Sample information browse | **√** | **√** | **√** |  |  | **√** | **√** |
|  | Detailed treatment information | **√** |  |  |  |  |  |  |
|  | Browsing by disease types | **√** |  | **√** |  |  |  | **√** |
|  | Browsing by biosample types & sources | **√** | **√** | **√** | **√** |  | **√** | **√** |
|  | UCSC genome browser link | **√** | **√** |  | **√** |  |  | **√** |
| **Quality control** | Mapping quality | **√** | **√** |  |  |  | **√** |  |
|  | TSS enrichment score | **√** | **√** | **√** |  |  |  |  |
|  | Fraction of reads in peaks | **√** | **√** | **√** |  |  | **√** |  |
|  | Quality control plots | **√** | **√** | **√** |  |  | **√** |  |
| **Region Annotation** | Nearest genes | **√** | **√** | **√** | **√** | **√** | **√** | **√** |
|  | Genomic features | **√** |  | **√** |  |  | **√** |  |
|  | Functional enrichment | **√** |  |  |  |  | **√** |  |
|  | Super-enhancer | **√** |  | **√** | **√** | **√** | **√** | **√** |
|  | Common SNPs | **√** |  | **√** |  | **√** | **√** | **√** |
|  | Risk SNPs | **√** |  | **√** |  | **√** | **√** | **√** |
|  | SNPs associated with blood disorders | **√** |  |  |  |  |  |  |
|  | eQTLs | **√** |  | **√** |  |  | **√** | **√** |
|  | LD SNPs | **√** |  | **√** |  |  |  |  |
| **Analysis functions** | TF footprint analysis | **√** |  |  |  |  |  |  |
|  | Differential footprinting between different datasets/conditions | **√** |  |  |  |  |  |  |
|  | CRC analysis | **√** |  |  |  |  | **√** | **√** |
| **Integrative Analysis** | Common & disease-specific regions | **√** |  |  |  |  |  |  |

*NOTE*: ^a^Blood cancer; ATAC-seq, the assay of transposase accessible chromatin with sequencing; DNase-Seq, Dnase I hypersensitive sites sequencing; H3K27ac, H3K27 acetylation; ChIP-seq, chromatin immunoprecipitation followed by sequencing; TSS, transcription start site; SNP, single nucleotide polymorphism; eQTL, expression quantitative trait locus; LD, linkage disequilibrium; TF, transcription fact­­or; CRC, core transcription regulatory circuitry.
